# Visual speed compensation of pitch-constrained blue-bottle flies under retinal-image velocity perturbation in a flight mill

**DOI:** 10.1101/707000

**Authors:** Shih-Jung Hsu, Bo Cheng

## Abstract

In the presence of wind or background image motion, flies are able to maintain a constant retinal-image velocity via regulating flight speed to the extent permitted by their locomotor capacity. Here we investigated the speed regulation of semi-tethered blue-bottle flies (*Calliphora vomitoria*) flying along an annular corridor in a magnetically levitated flight mill enclosed by two motorized cylindrical walls. We perturbed the flies’ retinal-image motion via spinning the cylindrical walls, generating bilaterally-averaged velocity perturbations from -0.3 to 0.3 m·s^-1^. Flies compensated retinal-image velocity perturbations by adjusting airspeed up to 20%, thereby maintaining a relatively constant retinal-image velocity. When the retinal-image velocity perturbation became greater than ∼0.1 m·s^-1^, the compensation weakened as airspeed plateaued, suggesting that flies were unable to further change airspeed. The compensation gain, i.e., the ratio of airspeed compensation and retinal-image velocity perturbation, depended on the spatial frequency of the grating patterns, being the largest at 12 m^-1^.

## INTRODUCTION

Flying insects rely heavily on retinal-image motion to control their flight (Borst et al., 2010; Srinivasan and Zhang, 2004). They often regulate flight speed by maintaining a constant retinal-image velocity (or optic flow) (David, 1982; Serres et al., 2008; Srinivasan et al., 1996; Srinivasan et al., 2000), which can be extracted over a broad range of image spatiotemporal frequencies (Fry et al., 2009). This differs from the classical optomotor turning response to rotating grating patterns (Borst et al., 2010), which is sensitive to image temporal frequency that depends on the image spatial frequency at a given retinal-image velocity.

Visual control of forward flight in insects is commonly studied using flight tunnels in free-flight (David, 1982; Fuller et al., 2014; Serres et al., 2008; Srinivasan et al., 1996) or tethered-flight settings (David, 1978; Lawson and Srinivasan, 2017). Recent efforts by the authors also demonstrated the potential of using flight mills to study forward flight in semi-tethered settings with controlled flight conditions (Hsu et al., 2019). While evidence has shown that free-flying insects in flight tunnels are able to maintain a constant retinal-image velocity in the presence of wind or background image motion (David, 1982; Srinivasan et al., 1996), it is unknown whether the same speed-regulation behavior also remains in semi-tethered flies in a flight mill.

Flying insects change their flight speed mainly through body-pitch dominated vectoring and magnitude modulation of cycle-averaged wing aerodynamic force, with a relatively small change in its direction relative to the body, a mechanism commonly described as the “helicopter model” (David, 1978; Muijres et al., 2014). However, it has been observed that insects can also change their flight speeds in the absence of any pitch maneuvers. For example, fruit flies can use a swimming-like paddling wing motion for thrust and speed modulation (Ristroph et al., 2011); and drone flies can produce significant airspeed changes, with negligible body pitch variations around 0°, at least for a short period of time (3-10 seconds) (Meng and Sun, 2016). As these relatively scarce observations suggest that insects are at least capable of relying only on the changes in wing motion to fly at different speeds, therefore do not have to conform to the helicopter model, it is unclear if they still modulate flight speed to maintain a constant retinal-image velocity with a constrained pitch, and to what extent can the changes of wing motion modulate speed before the locomotor limit is reached.

In this study, we used a motorized magnetically levitated (MAGLEV) flight mill, modified from those used in Hsu et al. (Hsu et al., 2019), to study the steady-state speed regulation in blue bottle flies (*Calliphora vomitoria*). We aimed to address two questions: 1) whether flies attempt to maintain a constant retinal-image velocity under retinal-image velocity perturbations when flying in the flight mill? similar to those observed in free flight, and 2) whether and to what extent can flies regulate flight speed using only the wing kinematic changes without body pitch change? In the experiments, the flies were constrained at a fixed body pitch (0°) in an annular corridor between two cylindrical walls displaying grating patterns, which generated retinal-image velocity perturbations via their co-rotations at the same angular velocity. We quantified the flies’ responses in terms of the changes in airspeeds and retinal-image velocities, as functions of retinal-image velocity perturbations.

## MATERIALS AND METHODS

### (a) Motorized magnetic levitated (MAGLEV) flight mill apparatus

A motorized MAGLEV flight mill (Fig. 1A), modified from that developed in Hsu and his coworkers (Hsu et al., 2019), was used in this study. The flight mill shaft, with three permanent magnets placed at its center, was levitated by two vertically aligned electromagnets. A proportional-integral-derivative (PID) controller generated pulse width modulation (PWM) signals to the electromagnets using the feedback of the permanent magnets’ vertical position measured by a linear Hall effect sensor. The flies, tethered to one end of the shaft, flew in an annular corridor of the flight mill enclosed by an inner (diameter 0.254 m) and an outer (diameter 0.349 m) cylindrical wall, which were centered via a vertical metal post and were driven by stepper motors via timing gears and pulleys. Both cylindrical walls displayed vertical grating patterns with one of the three spatial frequencies (SF): 24, 12 and ~0 (no pattern) m^−1^, i.e., the number of the grating cycles per meter. Note that all angular units in the paper were converted to linear units using the radius of the fly’s circular path at the center of the annular corridor, which was 0.146 m. The syn-directional and anti-directional rotations of the cylindrical walls (or the translations of the grating patterns) relative to the flight direction of the flies, created perturbations to the fly’s retinal-image velocity. We mounted two laser-and-photodiode pairs on the metal post with a 120° spacing (Fig. 1A), and we measured the average ground speed of the flies (*V*_*g*_, the forward velocity of the fly with respect to the fixed laboratory frame) based on the time lapses of the shaft passing two of the laser-and-photodiode pairs. A light source was mounted on the top of the entire device to provide consistent lighting (2877 ± 868 lux) to the annular corridor.

**Figure 1.**
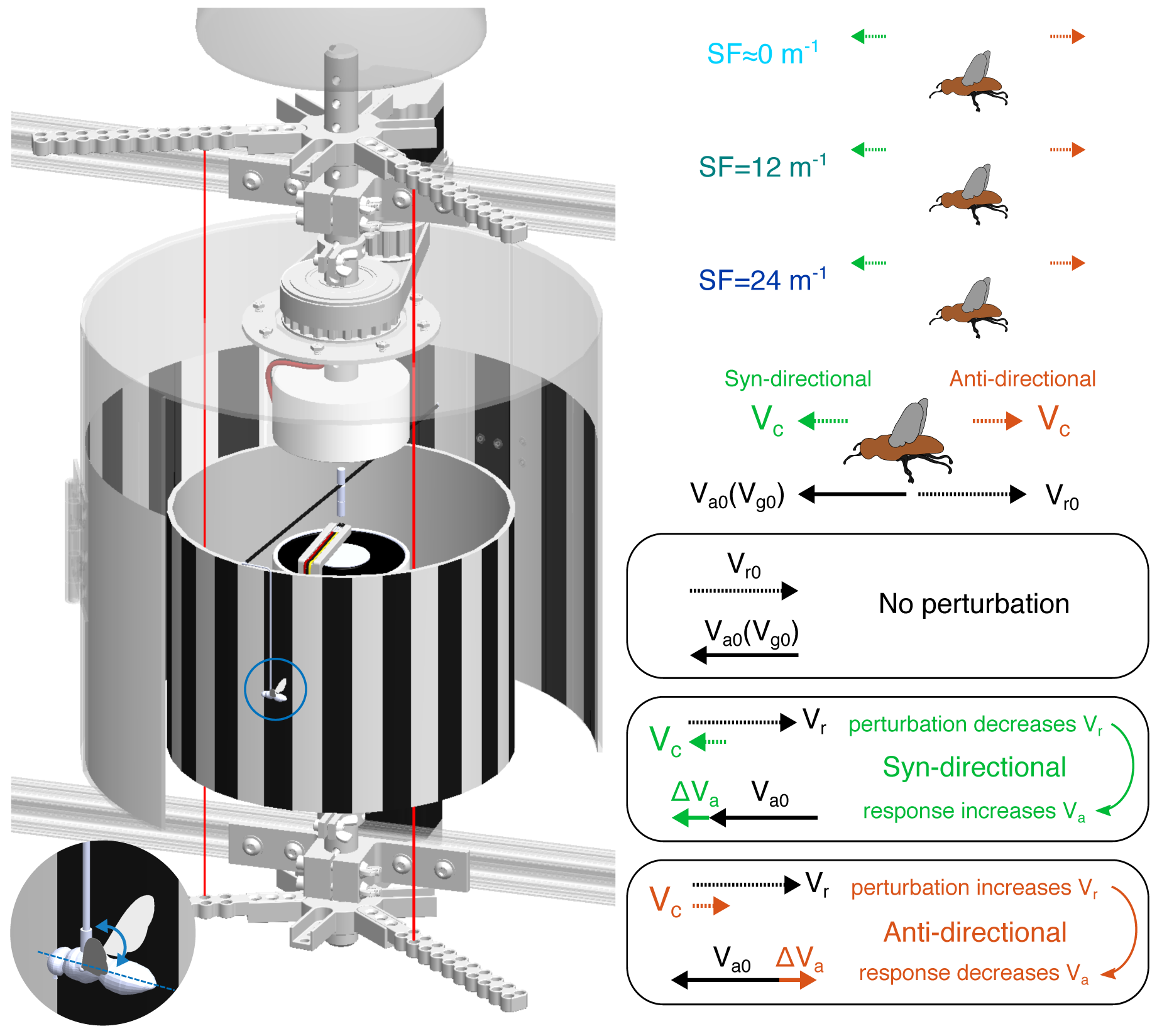
Experimental apparatus and methods. (A) The design of the MAGLEV flight mill system: a) timing gears, pulleys and stepper motors, b) permanent magnets and mill rotating shaft, c) Hall effect sensor, d) electromagnets, e) laser-and-photodiode pairs, f) outer and g) inner cylindrical walls with squared wave grating patterns. Magnified cutaway view: blue-bottle fly tethered at 0° body pitch with an angle pin. (B) Grating patterns with three spatial frequencies (SF=~0, 12 and 24 m^−1^) used in the experiments. (C) Retinal-image velocity perturbations (*V*_*c*_) are given syn-directionally (green dotted arrow) or anti-directionally (red dotted arrow). With no perturbation, the fly flies with an airspeed *V*_*a*0_ (same as ground speed *V*_*g*0_) generating an opposite retinal-image velocity *V*_*r*0_. Syn-directional (anti-directional) perturbation *V*_*c*_ decreases (increases) the retinal-image velocity *V*_*r*_, which is compensated by the fly via increasing (decreasing) its airspeed (Δ*V*_*a*_).

### (b) Animal preparation and experimental procedure

We used 4- to 8- days blue-bottle flies (*Calliphora vomitoria*) (N = 42, 32.9 ± 8.7 mg) for the experiments. Each fly was first cold anesthetized, while having its thorax glued orthogonally to a metal pin. The metal pin was then attached orthogonally to the flight mill shaft, therefore holding the fly at approximately 0° body pitch angle (Fig. 1A). After being introduced into the flight mill, the flies flew continuously in a clockwise direction at the middle of the annular corridor between the two cylindrical walls (Fig. 1A, also see Movie S1). Note that since the distance between the flies and the inner and outer walls were fixed for all experiments, the linear velocity of the grating patterns on the rotating cylindrical walls was equivalent to the image velocity or the optical flow introduced to a fly’s retina in our study. The retinal-image velocity perturbations were applied to each fly with a 0.03 m·s^−1^ interval following two sequences with opposite directions: 1) 0 → 0.3 → −0.3 → 0 m·s^−1^, and 2) 0 → −0.3 → 0.3 → 0 m·s^−1^. The results of one-way analysis of variance (ANOVA) test showed that for each SF group, the direction of the sequences has no significant effect on flies’ mean forward velocity (Table S2). After each change of perturbation velocity, a fly’s forward velocity was measured after 40 seconds of waiting period to ensure that the fly flew at near steady state. Note that in the experiment, individual flies were not distinguished.

As the two cylindrical walls spun, they induced air flow between them (i.e., Taylor-Couette flow (Taylor, 1923)). This wall-induced wind speed (*V*_*w*_) led to a difference between the ground speed (directly measured) and the airspeed of the flies, and therefore needs to be considered in the calculation of the latter analysis. Therefore, *V*_*w*_ was measured for each perturbation velocity without the presence of the fly by a hot-wire anemometer (405i, testo, Lenzkirch, Germany) and was found linearly dependent on the spinning speed (Fig. S1).

### (c) Data analyses and model selection

The responses of the flies to retinal-image velocity perturbations (*V*_*c*_) were measured by the changes in their bilaterally-averaged retinal-image velocity (Δ*V*_*r*_) and the changes in their airspeed (Δ*V*_*a*_). The bilaterally-averaged retinal-image velocity (*V*_*r*_) perceived by the fly, which was equivalent to the relative groundspeed between the fly and the cylindrical walls, was calculated by,

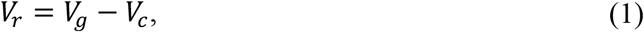

where *V*_*g*_ was the measured groundspeed and *V*_*c*_ was the bilaterally-averaged linear velocity of the grating patterns on inner and outer walls, which was equal to the product of the angular velocity of the cylindrical walls and the radius of the fly’s circular path at the center of the annular corridor. Note that in this study, we didn’t distinguish the retinal-image velocities between the two eyes of the flies, as both *V*_*r*_ and *V*_*c*_ were considered bilaterally averaged.

The fly’s airspeed (*V*_*a*_) was calculated based on the groundspeed (*V*_*g*_) and the wall-induced wind speed (*V*_*w*_),

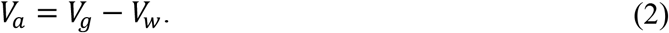

Note that we used the airspeed instead of the groundspeed in our analysis because the former better reflected the biomechanical efforts or the constraints of the flies in the compensation of the perturbation.

Assuming the existence of visual speed compensation, a fly would change its airspeed (or groundspeed) in the same direction of the retinal-image velocity perturbation, thereby keeping its retinal-image velocity near constant (or Δ*V*_*r*_ ≈ 0) (Fig. 1C). In addition, assuming that the changes of the airspeed (Δ*V*_*a*_) would saturate due to biomechanical constraints when the magnitude of the perturbation (*V*_*c*_) became sufficiently large, the flies would not be able to indefinitely increase or decrease their airspeed once it reached to an upper or lower bound, respectively. As a result, the Δ*V*_*r*_ would eventually increase (or decrease) with *V*_*c*_ as no further compensation would be possible.

Therefore, to model the flies’ response Δ*V*_*r*_ as a function of *V*_*c*_, we performed nonlinear regressions and model selection on two competing explanatory models developed assuming the existence or nonexistence of visual speed compensation, i.e., a linear function model 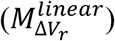 (assuming no visual compensation) and a cubic function model 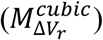 (assuming the existence of visual compensation). The cubic function was defined as,

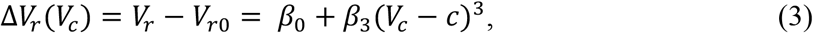

where *V*_*r*0_ was the measured mean retinal-image velocity of the flies without perturbation for each trial, *β*_0_ and *β*_3_ are the coefficients of polynomials, and *c* is the inflection point, *β*_0_, *β*_3_, and *c* were obtained from nonlinear regression.

To model the flies’ response Δ*V*_*a*_ as a function of *V*_*c*_, we also performed nonlinear regressions and model selection on two explanatory models: a linear function model (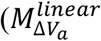, assuming no airspeed saturation) and a generalized logistic function model (Richards, 1959) 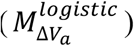 = (assuming airspeed saturation). The generalized logistic function is defined as:

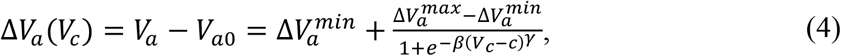

where *V*_*a*0_ was the measured mean airspeed of the flies without perturbation of each trial, 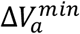 and 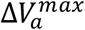 were the lower and upper asymptotic bounds, representing the minimal and maximum change in airspeed, respectively, *β* is the growth rate, *c* is where the maximum growth rate occurs and γ is the asymmetry coefficient (Richards, 1959). We also defined the compensation gain as the ratio between changes of airspeed and image-velocity perturbation, evaluated locally at *V*_*c*_ = 0:

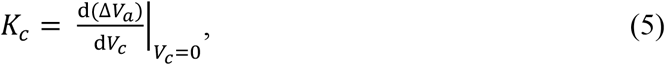

which was simply the slope for 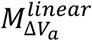 and the first derivative of 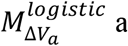 at *V*_*c*_ = 0.

We used the Levenberg-Marquardt algorithm (Elzhov et al., 2015) for the nonlinear regression to find the best fit for each model. Finally, the models were compared using Akaike information criterion (AIC) (Akaike, 1998), which evaluated the trade-off between the goodness-of-fit and the simplicity of the model.

## RESULTS AND DISCUSSION

42 blue-bottle flies (32.9 ± 8.7 mg) were used in the experiments, producing 2,228 measurements of flight speeds for the 21 cases of retinal-image perturbation (*V*_*c*_) ranging from −0.3 m·s^−1^ to 0.3 m·s^−1^ (see the sample size for each case in Table S1). The groundspeed (*V*_*g*_) at *V*_*c*_ = 0 was 0.44 ± 0.12 m·s^−1^ (Fig. S2) (no significant effect of SF on *V*_*g*_ at the p<.05 level [F_2, 201_ = 1.304, p = 0.274]).

When *V*_*c*_ were relatively small (−0.12 < *V*_*c*_ < 0.10 m·s^−1^, Fig. 2B), the flies well compensated the perturbations at SF = 12 m^−1^ by keeping retinal-image velocity approximately constant (Δ*V*_*r*_ ≈ 0) (Fig. 2A). The compensations existed but were significantly weaker at the higher (SF = 24 m^−1^) or lower (SF ~ 0 m^−1^) SF (Fig. 2A), both of which resulted in a near linear trend of Δ*V*_*r*_ in response to *V*_*c*_. When the magnitude of the perturbations became large, the compensation in SF = 12 m^−1^ group also begun to weaken as Δ*V*_*r*_ started to change linearly in the opposite direction of perturbation *V*_*c*_.

**Figure 2.**
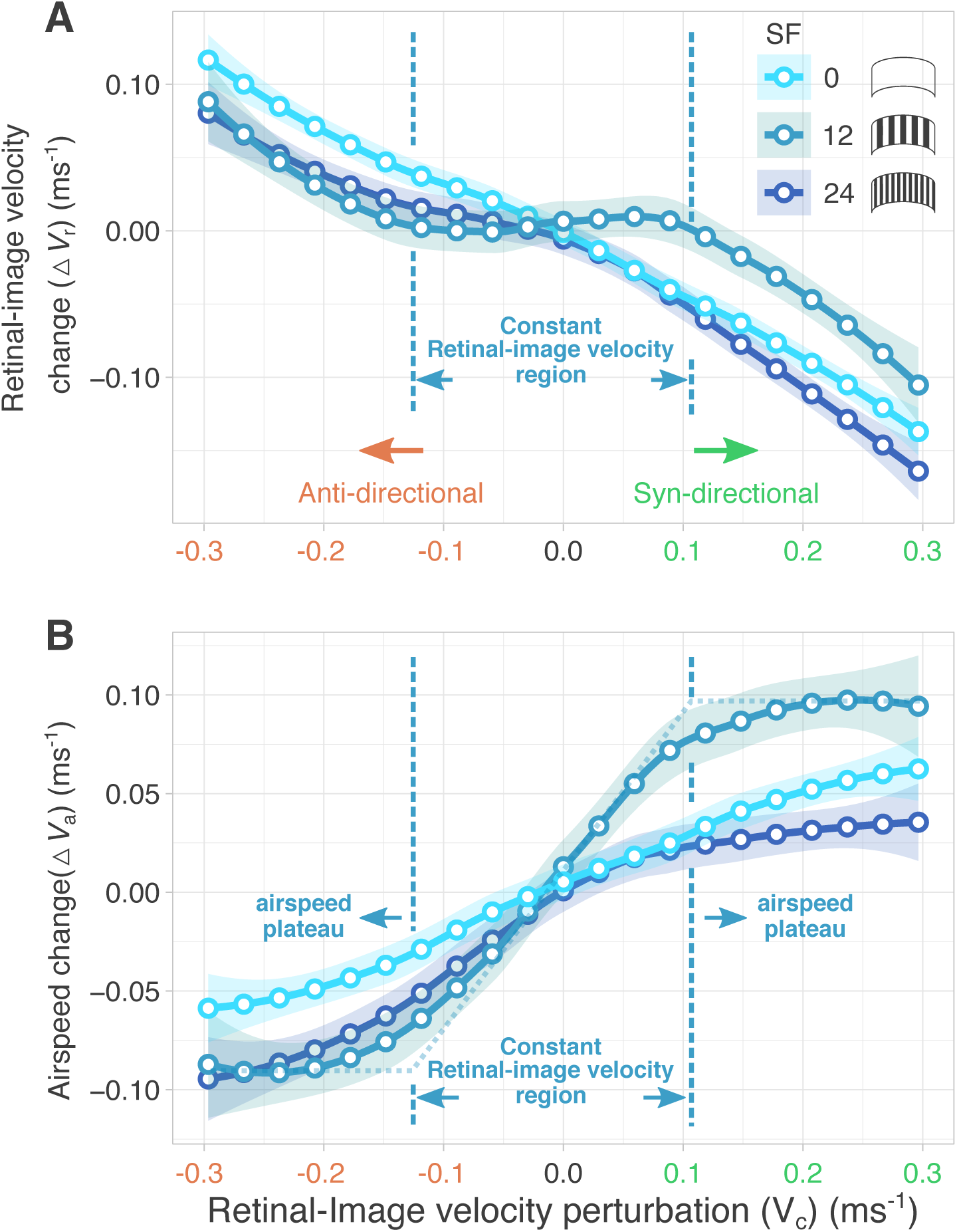
Flies’ responses under retinal-image velocity perturbation. (A) The change of flies’ retinal-image velocity (Δ*V*_*r*_) in response to the perturbation (*V*_*c*_). (B) The change of flies’ airspeed (Δ*V*_*a*_) in response to the perturbation (*V*_*c*_). Solid lines and shades represent LOESS regression mean and 95% confidence interval, respectively. Airspeed plateaued (at SF = 12) when retinal-image velocity perturbation is above 0.10 m·s^−1^ or below −0.12 m·s^−1^ (calculated by intersecting of the asymptote lines and the tangent line at the inflection point based on the data of SF=12 m^−1^. Teal dotted line in (B)). Between these two bounds (vertical dashed lines), blue bottle flies successfully maintained a constant retinal-image velocity. Horizontal green and red arrows indicate the syn-directional and the anti-directional image-velocity perturbations, respectively. Visual speed compensation corresponds to a syn-directional change of flies’ airspeed or groundspeed with respect to the perturbation, regardless of the perturbation is syn-directional or anti-directional to the flies absolute airspeed or groundspeed.

The observed compensation to maintain Δ*V*_*r*_ ≈ 0 was mainly the result of the changes in flies’ airspeed in response to the perturbation, as flies changed their airspeed in the same direction of the perturbation. This was confirmed by the monotonically trends of Δ*V*_*a*_ with *V*_*c*_ for all three SF cases (Fig. 2B). The compensation gain was significantly higher for SF = 12 m^−1^ (*K*_*c*_ = 0.868), compared with those of SF = 0 m^−1^ (*K*_*c*_ = 0.233) and SF = 24 m^−1^ (*K*_*c*_ = 0.337). The airspeed changes exhibited saturation under large perturbations, most apparently for SF = 12 and 24 m^−1^ (Fig. 2B). For SF = 24 m^−1^, the upper and lower bounds of Δ*V*_*a*_ were 0.034 ± 0.009 m·s^−1^ (mean ± 95% confidence interval (CI)) and −0.089 ± 0.010 m·s^−1^ (mean ± 95% CI), respectively. For SF=12 m^−1^, the upper and lower bounds of Δ*V*_*a*_ were 0.095 ± 0.010 m·s^−1^ and −0.091 ± 0.014 m·s^−1^ (mean ± 95% CI).

Consistent with the above observations, the model selection results (Table 1) showed that for SF = 12 m^−1^ the AIC favored models that assumed the existence of visual speed compensation and airspeed saturation, i.e., 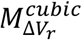 and 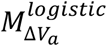, while for SF = 0 m^−1^ the AIC favored the models assuming no compensation and no airspeed saturation, i.e., 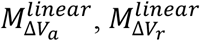. For SF = 24 m^−1^, AIC favored the model that assumed no visual speed compensation 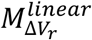 and the model that assumed the existence of airspeed saturation 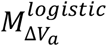.

**Table 1.** Model selection results based on Akaike’s information criterion (AIC). 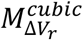 and 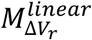 are the models for the retinal-image velocity response (Δ*V*_*r*_) assuming the existence and the nonexistence of visual speed compensation, respectively. 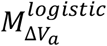 and 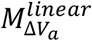 are models for the airspeed response (Δ*V*_*a*_) assuming the existence and the nonexistence of airspeed saturation, respectively. The selected models are highlighted in light gray. Nonlinear regression results are shown in Table S3.

| Retinal-image velocity change ($\Delta V_r$) | Model | | d.f. | Loglikely-hood | AIC | AIC weights |
| --- | --- | --- | --- | --- | --- | --- |
| SF=24 m<sup>-1</sup> | $M_{\Delta V_r}^{cubic}$ | $\Delta V_r = \beta_0 + \beta_3(V_c - c)^3$ | 3 | 798.05 | -1588.09 | <0.01 |
| | $M_{\Delta V_r}^{linear}$ | $V_r = \beta V_c + c$ | 2 | **807.39** | **-1608.79** | **>0.99** |
| SF=12 m<sup>-1</sup> | $M_{\Delta V_r}^{cubic}$ | $\Delta V_r = \beta_0 + \beta_3(V_c - c)^3$ | 3 | **560.03** | **-1112.07** | **>0.99** |
| | $M_{\Delta V_r}^{linear}$ | $V_r = \beta V_c + c$ | 2 | 550.12 | -1094.23 | <0.01 |
| SF=0 m<sup>-1</sup> | $M_{\Delta V_r}^{cubic}$ | $\Delta V_r = \beta_0 + \beta_3(V_c - c)^3$ | 3 | 1008.99 | -2009.98 | <0.01 |
| | $M_{\Delta V_r}^{linear}$ | $V_r = \beta V_c + c$ | 2 | **1048.20** | **-2090.40** | **>0.99** |
| Airspeed change ($\Delta V_a$) | Model | | d.f. | Loglikely-hood | AIC | AIC weights |
| SF=24 m<sup>-1</sup> | $M_{\Delta V_a}^{logistic}$ | $\Delta V_a = \Delta V_a^{min} + \frac{\Delta V_a^{max} - \Delta V_a^{min}}{1 + e^{-\beta(V_c - c)^\gamma}}$ | 5 | **820.98** | **-1629.95** | **>0.99** |
| | $M_{\Delta V_a}^{linear}$ | $V_a = \beta V_c + c$ | 2 | 534.07 | -1615.04 | <0.01 |
| SF=12 m<sup>-1</sup> | $M_{\Delta V_a}^{logistic}$ | $\Delta V_a = \Delta V_a^{min} + \frac{\Delta V_a^{max} - \Delta V_a^{min}}{1 + e^{-\beta(V_c - c)^\gamma}}$ | 5 | **562.41** | **-1112.82** | **>0.99** |
| | $M_{\Delta V_a}^{linear}$ | $V_a = \beta V_c + c$ | 2 | 550.29 | -1094.58 | <0.01 |
| SF=0 m<sup>-1</sup> | $M_{\Delta V_a}^{logistic}$ | $\Delta V_a = \Delta V_a^{min} + \frac{\Delta V_a^{max} - \Delta V_a^{min}}{1 + e^{-\beta(V_c - c)^\gamma}}$ | 5 | 1050.81 | -2089.62 | 0.27 |
| | $M_{\Delta V_a}^{linear}$ | $V_a = \beta V_c + c$ | 2 | **1048.82** | **-2091.63** | **0.73** |

In this work, we studied the responses of blue-bottle flies (*Calliphora vomitoria*) subjected to retinal-image velocity perturbation while flying in the motorized MAGLEV flight mill with a constrained pitch. Similar to those observed in free-flight conditions (David, 1982; Serres et al., 2008; Srinivasan et al., 1996; Srinivasan et al., 2000), the flies appeared to compensate retinal-image velocity perturbations by changing their airspeed (or groundspeed) syn-directionally with the perturbation. The compensation was the strongest for perturbations between −0.12 m·s^−1^ and 0.10 m·s^−1^ at SF = 12 m^−1^, corresponding to a compensation gain at 0.868, significantly higher than those for SF = ~0 and 24 m^−1^. The observed dependency on image spatial frequency in the speed compensation responses was similar to those identified in the free-flight acceleration responses of fruit flies (*Drosophila melanogaster*) (Fry et al., 2009), which showed that the response strength was among the strongest at SF of 12 m^−1^, but reduced substantially at 24 m^−1^. Nevertheless, compared with classical optomotor turning response (Borst et al., 2010), it is expected that the dependency on spatial frequency in speed compensation or acceleration responses is relatively weak, because the retinal-image velocity can be extracted robustly over a broad range of image spatiotemporal frequencies (Fry et al., 2009), while the optomotor turning response is proportional to image temporal frequency that depends on image spatial frequency at a fixed image velocity (Borst et al., 2010).

When the magnitude of the retinal-image velocity perturbation became large, speed compensation weakened as the changes in airspeed plateaued, as flies were unable to adjust their airspeed beyond approximately ±0.1 m·s^−1^ (or +22% and −21% compared to the average airspeed at *V*_*c*_ = 0 under SF = 12 m^−1^). This was possibly due to a compound of biomechanical constraints, including the constrained pitch and their force-vectoring ability via changes of wing motion (Hsu et al., 2019). Note that the current studies were limited to the steady-state responses, while flies are likely capable of large transient modulation of wing motion during rapid aerodynamic maneuvers and flight stabilization (Ristroph et al., 2010). It can be also speculated that with a free body pitch, e.g., by tethering a fly to the flight mill via a micro bearing, the successful compensation region (Fig. 2B) can be further expanded to larger image-velocity perturbations.

In conclusion, to address the two motivating questions of this study, our results showed that 1) when flying in the MAGLEV flight mill, flies did regulate their flight speed to maintain a constant retinal-image velocity under image velocity perturbations, similar to those observed in free flight, and 2) flies were capable of adjusting flight speed using only the wing kinematic changes during steady-state flight, however the amount of adjustment was relatively small (less than 0.1 m/s).

## ACKNOWLEDGEMENTS

We thank Elizabeth Seber and Ciera McFarland for the assistance with the experiments.

## AUTHOR CONTRIBUTIONS

S.J.H. and B.C. contributed to the design of the experiments, data analysis, interpretation of the findings and the manuscript preparation. S.J.H. prepared and conducted the experiments.

## SUPPLEMENTARY INFORMATION

Supplementary information available online at http://jeb.glologists.org/lookup/doi/xxx/xxx.supplemental

