## Supplementary Materials for "Visual speed compensation of pitch-constrained blue-bottle flies under retinal-image velocity perturbation in a flight mill"

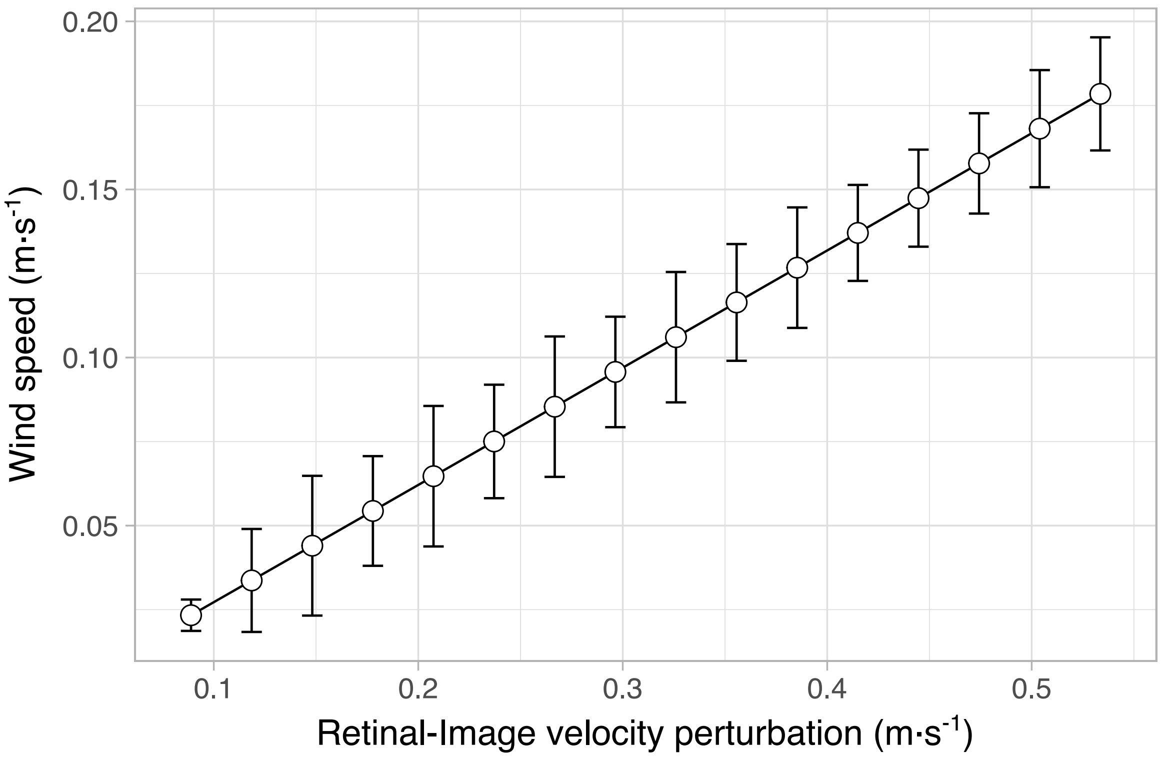


Figure S1. Induced wind speed (Taylor–Couette flow) at the center of the annular corridor between the two cylindrical walls as a function of retinal-image velocity perturbation (or cylinder spinning speed). Wind speeds were calibrated with an anemometer with two cylinders spinning simultaneously at the same speed in the same direction. The anemometer was placed at three separate locations along the flies' flight path. With each velocity change, we waited 120 seconds between the measurements, and total 60 measurement samples were collected at each velocity at 0.5 Hz sampling rate. As the cylinder velocity increased, the wind speed in the corridor increased linearly.

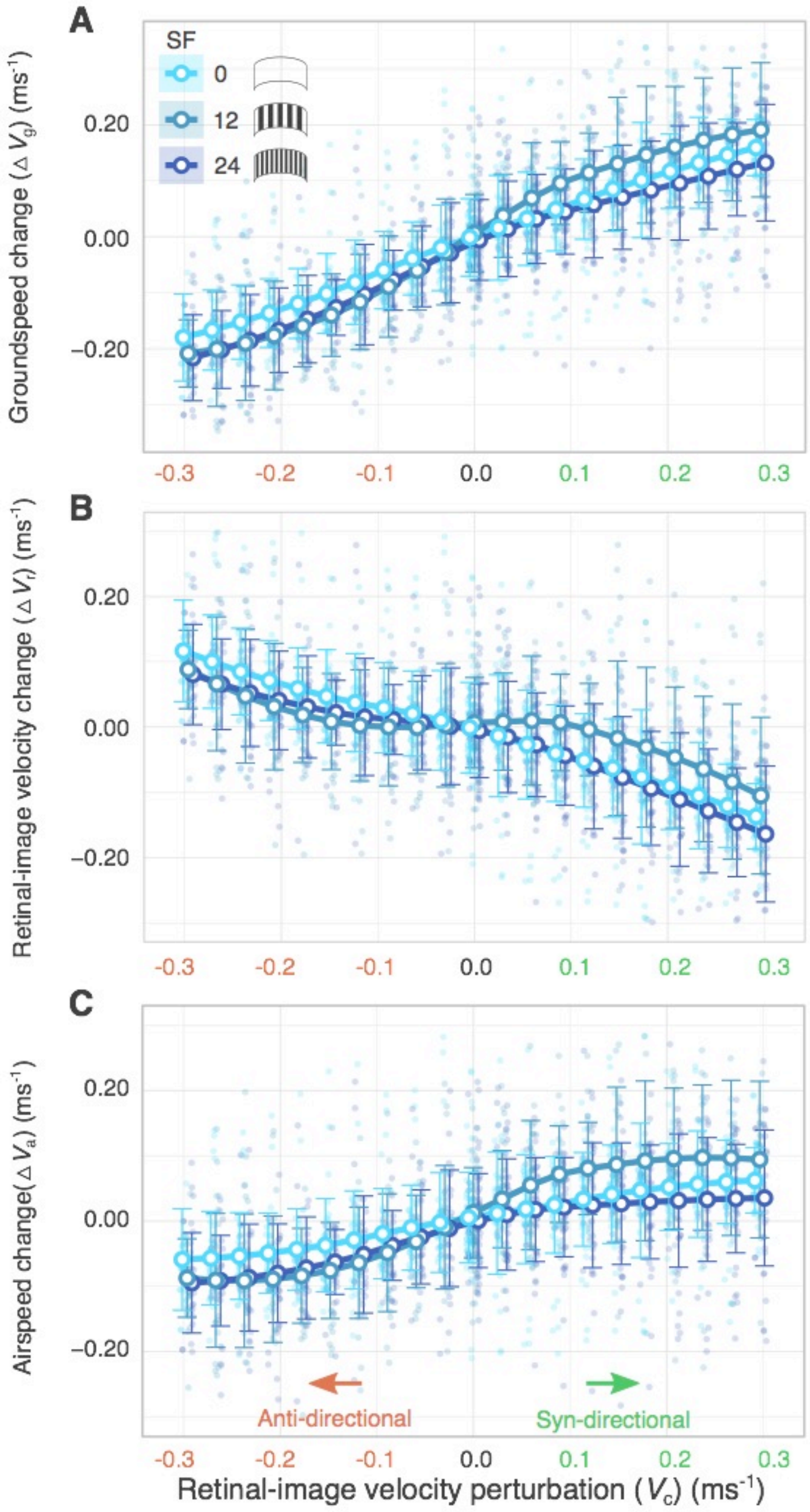


Figure S2. The flies' responses under retinal image velocity perturbation shown with the raw data. (A) Groundspeed change, (B) retinal-image velocity change and (C) airspeed change in response to retinal-image velocity perturbation ( $V_c$ ). The results for three spatial frequency (SF) groups are plotted in dark blue (SF=24), teal (SF=12) and light blue (SF=0) colours. Raw data, regression lines and error bars are slightly shifted along x-axis for better visibility. The raw measurement data are shown in transparent background points, together with the LOESS regression line (solid lines) and the standard deviations (error bars).

16

Table S1. Detailed sample size arranged according to spatial frequency (SF), direction of sequence (seq) and the given retinal-image velocity perturbation ($V_c$). Total sample size in each SF group: 761 (SF=24), 587 (SF=12), and 880 (SF=~0).

| Retinal-image velocity perturbation $V_c$ (m·s$^{-1}$) | | -0.3 | -0.27 | -0.24 | -0.21 | -0.18 | -0.15 | -0.12 | -0.09 | -0.06 | -0.03 | 0 |
|---|---|---|---|---|---|---|---|---|---|---|---|---|
| SF=24 | seq: 0→0.3→-0.3→0 | 16 | 17 | 17 | 17 | 17 | 17 | 17 | 17 | 17 | 17 | 39 |
| | seq: 0→-0.3→0.3→0 | 12 | 13 | 13 | 13 | 13 | 13 | 13 | 13 | 13 | 13 | 31 |
| | total sample size | 28 | 30 | 30 | 30 | 30 | 30 | 30 | 30 | 30 | 30 | 70 |
| SF=12 | seq: 0→0.3→-0.3→0 | 12 | 12 | 12 | 12 | 12 | 13 | 13 | 13 | 13 | 13 | 28 |
| | seq: 0→-0.3→0.3→0 | 12 | 12 | 12 | 12 | 12 | 12 | 12 | 12 | 12 | 12 | 26 |
| | total sample size | 24 | 24 | 24 | 24 | 24 | 25 | 25 | 25 | 25 | 25 | 54 |
| SF=~0 | seq: 0→0.3→-0.3→0 | 18 | 18 | 18 | 18 | 18 | 18 | 18 | 18 | 18 | 18 | 40 |
| | seq: 0→-0.3→0.3→0 | 18 | 18 | 18 | 18 | 18 | 18 | 18 | 18 | 18 | 18 | 40 |
| | total sample size | 36 | 36 | 36 | 36 | 36 | 36 | 36 | 36 | 36 | 36 | 80 |
| Retinal-image velocity perturbation $V_c$ (m·s$^{-1}$) | | 0.3 | 0.27 | 0.24 | 0.21 | 0.18 | 0.15 | 0.12 | 0.09 | 0.06 | 0.03 | |
| SF=24 | seq: 0→0.3→-0.3→0 | 19 | 21 | 21 | 21 | 21 | 22 | 22 | 22 | 22 | 22 | |
| | seq: 0→-0.3→0.3→0 | 18 | 18 | 18 | 18 | 18 | 18 | 18 | 18 | 18 | 18 | |
| | total sample size | 37 | 39 | 39 | 39 | 39 | 40 | 40 | 40 | 40 | 40 | |
| SF=12 | seq: 0→0.3→-0.3→0 | 14 | 14 | 15 | 15 | 15 | 15 | 15 | 15 | 15 | 15 | |
| | seq: 0→-0.3→0.3→0 | 14 | 14 | 14 | 14 | 14 | 14 | 14 | 14 | 14 | 14 | |
| | total sample size | 28 | 28 | 29 | 29 | 29 | 29 | 29 | 29 | 29 | 29 | |
| SF=~0 | seq: 0→0.3→-0.3→0 | 22 | 22 | 22 | 22 | 22 | 22 | 22 | 22 | 22 | 22 | |
| | seq: 0→-0.3→0.3→0 | 22 | 22 | 22 | 22 | 22 | 22 | 22 | 22 | 22 | 22 | |
| | total sample size | 44 | 44 | 44 | 44 | 44 | 44 | 44 | 44 | 44 | 44 | |

Table S2. One-way analysis of variance (ANOVA) test on the significance of velocity sequence direction on blue-bottle flies' mean groundspeed ($V_g$) for each image-perturbation. $H_0$: direction of the sequences has no effect on blue-bottle flies' mean groundspeed. $H_a$: direction of the sequences has an effect on blue-bottle flies' mean groundspeed. The level of significance was p=0.05. The test was evaluated with the function aov in R stats package. Results showed that for 59 out of 60 cases (3 spatial frequencies × 20 retinal-image perturbations), there were no significant differences in the mean groundspeeds between the two directions of the sequence groups, except SF=24 at $V_c$=-0.03 m·s$^{-1}$ (indicated by *).

| Retinal-image velocity perturbation $V_c$ (m·s$^{-1}$) | | -0.3 | -0.27 | -0.24 | -0.21 | -0.18 | -0.15 | -0.12 | -0.09 | -0.06 | -0.03 | 0 |
|---|---|---|---|---|---|---|---|---|---|---|---|---|
| SF=24 | F value | N/A | 1.776 | 3.012 | 0.243 | 0.293 | 0.340 | 0.133 | 0.259 | 0.674 | 5.267 | 2.885 |
| | p value | N/A | 0.193 | 0.094 | 0.626 | 0.593 | 0.565 | 0.719 | 0.615 | 0.419 | 0.03* | 0.100 |
| SF=12 | F value | N/A | 0.080 | 0.014 | 0.490 | 0.392 | 0.369 | 0.050 | 0.085 | 0.472 | 0.737 | 0.096 |
| | p value | N/A | 0.779 | 0.907 | 0.491 | 0.538 | 0.549 | 0.825 | 0.773 | 0.499 | 0.399 | 0.759 |
| SF=0 | F value | N/A | 1.316 | 1.127 | 0.745 | 0.682 | 1.708 | 1.177 | 1.013 | 1.167 | 1.233 | 2.163 |
| | p value | N/A | 0.286 | 0.353 | 0.533 | 0.570 | 0.185 | 0.334 | 0.400 | 0.338 | 0.314 | 0.112 |
| Retinal-image velocity perturbation $V_c$ (m·s$^{-1}$) | | 0.3 | 0.27 | 0.24 | 0.21 | 0.18 | 0.15 | 0.12 | 0.09 | 0.06 | 0.03 | 0 |
| SF=24 | F value | N/A | 1.642 | 0.326 | 1.662 | 1.148 | 0.486 | 1.684 | 0.375 | 0.034 | 0.174 | 0.392 |
| | p value | N/A | 0.208 | 0.572 | 0.205 | 0.291 | 0.49 | 0.202 | 0.544 | 0.855 | 0.679 | 0.535 |
| SF=12 | F value | N/A | 3.661 | 2.582 | 0.289 | 0.796 | 0.002 | 0.34 | 0.009 | 0.211 | 0.676 | 0.637 |
| | p value | N/A | 0.067 | 0.12 | 0.595 | 0.38 | 0.964 | 0.565 | 0.927 | 0.65 | 0.418 | 0.432 |
| SF=0 | F value | N/A | 0.445 | 0.755 | 0.325 | 1.872 | 0.945 | 1.150 | 1.265 | 0.883 | 2.032 | 0.964 |
| | p value | N/A | 0.722 | 0.526 | 0.807 | 0.150 | 0.428 | 0.341 | 0.300 | 0.458 | 0.125 | 0.419 |

Table S3. Nonlinear regression results. Significance levels: p<0.01**, p<0.05*.

| $M_{\Delta V_a}^{logistic}$ | Formula: $\Delta V_a = \Delta V_a^{min} + \frac{\Delta V_a^{max} - \Delta V_a^{min}}{1+e^{-\beta (V_c - c)^{\gamma}}}$ | | | |
|---|---|---|---|---|
| SF=24 | Estimate | Std. Error | t-value | Pr(>\|t\|) |
| $\Delta V_a^{min}$ | -0.10 | 0.03 | -2.94 | <0.01** |
| $\Delta V_a^{max}$ | 0.03 | 0.01 | 4.96 | <0.01** |
| $\beta$ | 22.55 | 20.26 | 1.11 | 0.27 |
| $\gamma$ | 0.42 | 0.92 | 0.46 | 0.65 |
| $c$ | -0.01 | 0.12 | -0.08 | 0.94 |

Residual standard error: 0.08254 on 756 d.o.f.

| $M_{\Delta V_a}^{linear}$ | Formula: $V_a = \beta V_c + c$ | | | |
|---|---|---|---|---|
| SF=24 | Estimate | Std. Error | t-value | Pr(>\|t\|) |
| $c$ | -0.02 | <0.01 | -5.37 | <0.01** |
| $\beta$ | 0.25 | 0.02 | 14.50 | <0.01** |

Residual standard error: 0.08352 on 759 d.o.f.

| $M_{\Delta V_a}^{logistic}$ | Formula: $\Delta V_a = \Delta V_a^{min} + \frac{\Delta V_a^{max} - \Delta V_a^{min}}{1+e^{-\beta (V_c - c)^{\gamma}}}$ | | | |
|---|---|---|---|---|
| SF=12 | Estimate | Std. Error | t-value | Pr(>\|t\|) |
| $\Delta V_a^{min}$ | -0.09 | 0.01 | -6.20 | <0.01** |
| $\Delta V_a^{max}$ | 0.10 | 0.01 | 9.46 | <0.01** |
| $\beta$ | 19.17 | 11.49 | 1.67 | 0.10 |
| $\gamma$ | 0.94 | 1.64 | 0.57 | 0.57 |
| $c$ | -0.01 | 0.12 | -0.07 | 0.95 |

Residual standard error: 0.09322 on 582 d.o.f.

| $M_{\Delta V_a}^{linear}$ | Formula: $V_a = \beta V_c + c$ | | | |
|---|---|---|---|---|
| SF=12 | Estimate | Std. Error | t-value | Pr(>\|t\|) |
| $c$ | 0.01 | 0.00 | 1.73 | 0.08 |
| $\beta$ | 0.41 | 0.02 | 18.10 | <0.01** |

Residual standard error: 0.09492 on 585 d.o.f.

| $M_{\Delta V_a}^{logistic}$ | Formula: $\Delta V_a = \Delta V_a^{min} + \frac{\Delta V_a^{max} - \Delta V_a^{min}}{1+e^{-\beta (V_c - c)^{\gamma}}}$ | | | |
|---|---|---|---|---|
| SF=0 | Estimate | Std. Error | t-value | Pr(>\|t\|) |
| $\Delta V_a^{min}$ | -0.07 | 0.05 | -1.42 | 0.16 |
| $\Delta V_a^{max}$ | 0.07 | 0.03 | 2.15 | 0.032* |
| $\beta$ | 7.52 | 9.70 | 0.78 | 0.44 |
| $\gamma$ | 1.77 | 11.38 | 0.16 | 0.88 |
| $c$ | -0.10 | 1.04 | -0.10 | 0.92 |

Residual standard error: 0.07352 on 875 d.o.f.

| $M_{\Delta V_a}^{linear}$ | Formula: $V_a = \beta V_c + c$ | | | |
|---|---|---|---|---|
| SF=0 | Estimate | Std. Error | t-value | Pr(>\|t\|) |
| $c$ | 0.00 | <0.01 | 1.10 | 0.27 |
| $\beta$ | 0.23 | 0.01 | 16.43 | <0.01** |

Residual standard error: 0.07356 on 878 d.o.f.

| $M^{cubic}_{\Delta V_r}$ | Formula: $\Delta V_r = \beta_0 + \beta_3 (V_c - c)^3$ | | | |
|---|---|---|---|---|
| SF=24 | Estimate | Std. Error | t-value | Pr(>\|t\|) |
| $\beta_0$ | -0.01 | 0.00 | -3.24 | <0.01** |
| $\beta_3$ | -5.54 | 0.32 | -17.58 | <0.01** |
| c | -0.03 | 0.01 | -3.67 | <0.01** |

Residual standard error: 0.08495 on 758 d.o.f.

| $M^{linear}_{\Delta V_r}$ | Formula: $V_a = \beta V_c + c$ | | | |
|---|---|---|---|---|
| SF=24 | Estimate | Std. Error | t-value | Pr(>\|t\|) |
| $c$ | -0.03 | 0.00 | -8.51 | <0.01** |
| $\beta$ | -0.38 | 0.02 | -21.60 | <0.01** |

Residual standard error: 0.08386 on 759 d.o.f.

| $M^{cubic}_{\Delta V_r}$ | Formula: $\Delta V_r = \beta_0 + \beta_3 (V_c - c)^3$ | | | |
|---|---|---|---|---|
| SF=12 | Estimate | Std. Error | t-value | Pr(>\|t\|) |
| $\beta_0$ | 0.00 | 0.01 | 0.33 | 0.74 |
| $\beta_3$ | -3.83 | 0.37 | -10.48 | <0.01** |
| c | -0.01 | 0.01 | -1.23 | 0.22 |

Residual standard error: 0.09344 on 584 d.o.f.

| $M^{linear}_{\Delta V_r}$ | Formula: $V_a = \beta V_c + c$ | | | |
|---|---|---|---|---|
| SF=12 | Estimate | Std. Error | t-value | Pr(>\|t\|) |
| $c$ | 0.00 | 0.00 | -0.73 | 0.46 |
| $\beta$ | -0.23 | 0.02 | -9.99 | <0.01** |

Residual standard error: 0.09495 on 585 d.o.f.

| $M^{cubic}_{\Delta V_r}$ | Formula: $\Delta V_r = \beta_0 + \beta_3 (V_c - c)^3$ | | | |
|---|---|---|---|---|
| SF=0 | Estimate | Std. Error | t-value | Pr(>\|t\|) |
| $\beta_0$ | -0.01 | 0.00 | -1.67 | 0.09 |
| $\beta_3$ | -5.99 | 0.24 | -25.23 | <0.01** |
| c | 0.00 | 0.01 | -0.82 | 0.42 |

Residual standard error: 0.07701 on 877 d.o.f.

| $M^{linear}_{\Delta V_r}$ | Formula: $V_a = \beta V_c + c$ | | | |
|---|---|---|---|---|
| SF=0 | Estimate | Std. Error | t-value | Pr(>\|t\|) |
| $c$ | -0.01 | 0.00 | -2.81 | <0.01** |
| $\beta$ | -0.40 | 0.01 | -28.20 | <0.01** |

Residual standard error: 0.07361 on 878 d.o.f.

Movie S1. Sample video of MAGLEV flight mill experiments: filghtmill.mp4 (http://figshare.com/articles/flightmill_mp4/8945276)
